# An ingestible, battery-free, tissue-adhering robotic interface for non-invasive and chronic electrostimulation of the gut

**DOI:** 10.1101/2024.04.25.591220

**Authors:** Kewang Nan, Kiwan Wong, Dengfeng Li, Binbin Ying, James C McRae, Vivian R Feig, Shubing Wang, Kuanming Yao, Jingkun Zhou, Jian Li, Joshua Jenkins, Keiko Ishida, Johannes Kuosmanen, Wiam Abdalla Mohammed Madani, Alison Hayward, Khalil Ramadi, Xinge Yu, Giovanni Traverso

**Affiliations:** College of Pharmaceutical Sciences, Zhejiang University, Hangzhou, China; Department of Mechanical Engineering, Massachusetts Institute of Technology, Cambridge, MA, USA; Division of Gastroenterology, Hepatology and Endoscopy, Brigham and Women’s Hospital, Harvard Medical School, Boston, MA, USA; Department of Biomedical Engineering, City University of Hong Kong, Hong Kong SAR, China; Hong Kong Center for Cerebro-Cardiovascular Health Engineering (COCHE), Hong Kong SAR, China; David H. Koch Institute for Integrative Cancer Research, Massachusetts Institute of Technology, Cambridge, MA, USA; Division of Comparative Medicine, Massachusetts Institute of Technology, Cambridge, MA, USA; Division of Engineering, New York University Abu Dhabi, Abu Dhabi, UAE; Tandon School of Engineering, New York University, New York, NY, USA

## Abstract

Ingestible electronics have the capacity to transform our ability to effectively diagnose and potentially treat a broad set of conditions. Current applications could be significantly enhanced by addressing poor electrode-tissue contact, lack of navigation, short dwell time, and limited battery life. Here we report the development of an ingestible, battery-free, and tissue-adhering robotic interface (IngRI) for non-invasive and chronic electrostimulation of the gut, which addresses challenges associated with contact, navigation, retention, and powering (C-N-R-P) faced by existing ingestibles. We show that near-field inductive coupling operating near 13.56 MHz was sufficient to power and modulate the IngRI to deliver therapeutically relevant electrostimulation, which can be further enhanced by a bio-inspired, hydrogel-enabled adhesive interface. In swine models, we demonstrated the electrical interaction of IngRI with the gastric mucosa by recording conductive signaling from the subcutaneous space. We further observed changes in plasma ghrelin levels, the “hunger hormone,” while IngRI was activated *in vivo*, demonstrating its clinical potential in regulating appetite and treating other endocrine conditions. The results of this study suggest that concepts inspired by soft and wireless skin-interfacing electronic devices can be applied to ingestible electronics with potential clinical applications for evaluating and treating gastrointestinal conditions.

## INTRODUCTION

Similar to brain, heart, and vagus nerve, the gastrointestinal (GI) tract consists of cells that exhibit electrical excitability and therefore can be modulated by electrostimulation (**Figure 1A**). In the GI tract, where the neural, endocrine, and immune systems intersect, electrostimulation has been shown to exert therapeutic effects against nausea^1^, vomiting^2^, gastric dysmotility^3^, obesity^4^, and diabetes^5^. In addition to treating GI disorders, neuromodulation has the potential to treat neurodegenerative diseases such as Alzheimer’s and Parkinson’s^6,7^, as it has been shown that the central and peripheral nervous systems are interrelated through the microbiota-gut-brain axis^8–10^.

**Figure 1:**
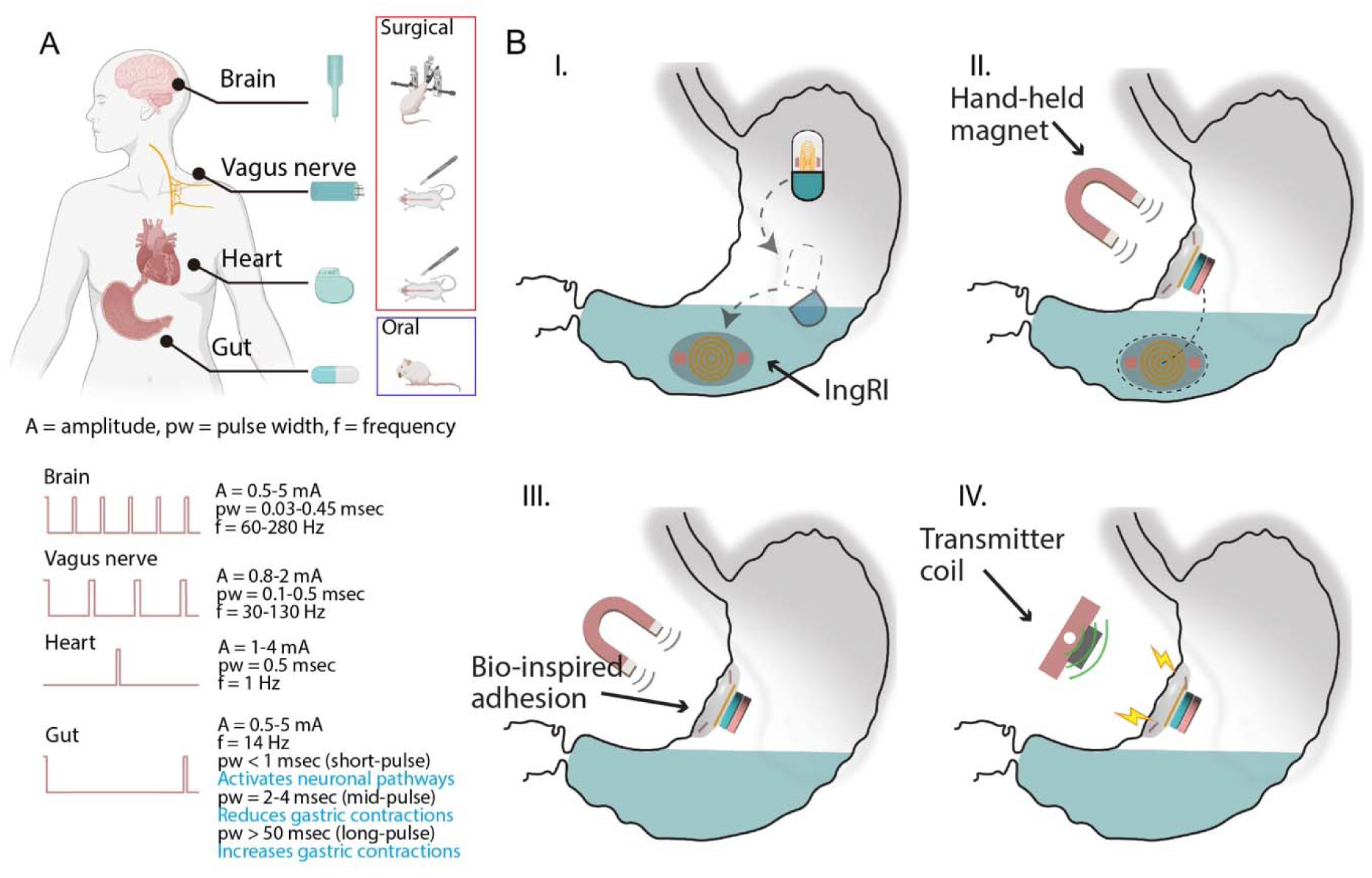
**The ingestible, battery-free, and tissue-adhering robotic interface (IngRI),** an electrostimulation device for wireless, long-term electrostimulation in the gut. (A) Therapeutically relevant electrostimulation for organs and tissues that exhibit electrical excitability and their typical delivery methods. (B) The mechanism of action of IngRI: (I) The gelatin capsule dissolves and releases the folded IngRI in the gut. (II) The IngRI navigates to the desired gastric location aided by an external hand-held magnet. (III) The IngRI adheres onto the gastric mucosa with bio-inspired hydrogel adhesion and activates conformal electrode-tissue contact. (IV) The IngRI delivers programmable electrical pulses via near-field inductive coupling.

From a device implementation perspective, the GI tract is appealing as it is among the few internal organs that can be accessed noninvasively via the oral route. Implantable electrostimulation devices for the brain^11^, vagus nerve^12^, and heart^13^ require surgical access, which may impede their widespread acceptance and accessibility to patients. Stimulators targeting the GI tract, on the other hand, can be designed to support self-administration in non-clinical settings^14,15^ (**Figure 1A**). However, existing ingestible devices for electrostimulation (IDEs) cannot yet achieve performance on par with their surgically implanted counterparts owing to issues associated with contact, navigation, retention, and powering (C-N-R-P):

### Contact

Most IDEs experience poor electrode-tissue contact due to random gastric positioning following ingestion, as well as mechanical mismatch between rigid capsules^14,16^ and soft gastric tissues, leading to highly variable and non-therapeutic interactions with the gastric wall^17^.

### Navigation

Most IDEs lack the autonomous locomotive ability to position their electrode leads precisely inside the gastric lumen without endoscopic interventions. This location has been shown to influence the effectiveness of electrostimulation^18,19^.

### Retention

Most IDEs cannot be retained in the gut for more than several hours^14,16^, which is in sharp contrast to surgically implanted gastric stimulators that can remain physically attached for years^20^. The recently reported use of barbed needles^17^ or expandable structures^21,22^ may extend the gastric retention time though may still experience poor electrode-tissue contact and additional risks such as tissue disruption that would have to be mitigated.

### Powering

Most IDEs have finite battery lives supplied by low-power density, non-rechargeable button-type batteries and have limited control over the electrical parameters of the output waveform, such as amplitude, pulse width, and frequency, which play central roles in determining the outcomes of electrostimulation (**Figure 1A**). These limitations stem from the highly constrained volume budgets when designing ingestible electronics, in which more than 50% of the space is currently consumed by batteries^14,16^. Accidental exposure of battery components to a corrosive gastric environment also increases the risk of obstruction, poisoning, and explosion^23^.

Recognizing many similarities shared by the gastric mucosa and skin^15^, we hypothesize that a thin, flexible, and skin-like device architecture could provide an enhanced and durable electronic tissue interface free of sutures, with wireless powering and modulation capabilities to eliminate battery needs^24–27^. However, additional design considerations are imposed by this unique gastric environment. First, the gastric mucosa is not as accessible as the skin, which complicates device deployment and navigation. Second, the gastric mucosa is surrounded by gastric fluid, contains creases and folds, and is subject to constant peristaltic mechanical motion, all of which are detrimental to the quality and durability of electrode-tissue contact. Third, near-field inductive coupling has not yet been shown to provide sufficient powering and modulating capabilities to devices in the GI tract because of the signal attenuation caused by the surrounding abdominal tissues.

Here, we describe the development of an **ingestible, battery-free, and tissue-adhering robotic interface (IngRI)** that fully addresses C-N-R-P issues. Comprised of photolithographically defined thin-film circuits embedded in a sub-millimeter thick elastomer substrate, the IngRI mechanically conforms to the gastric mucosa and forms robust electrode-tissue contact after being released from a commercial gelatin capsule (**Figure 1B, step I**). A small on-board permanent magnet allows remote orientation and navigation of the IngRI to a designated gastric position assisted by an external handheld magnet (**Figure 1B, step II**). A bio-inspired, hydrogel/elastomer hybrid structure was then activated within minutes to establish long-term retention in the gastric mucosa for at least 48 h (**Figure 1B, step III**). Finally, programmable electrical pulses were delivered using near-field inductive coupling operated near 13.56 MHz (**Figure 1B, step IV**). Using *in vivo* swine models, we showed that IngRI allowed wireless delivery of therapeutically relevant electrostimulation through a 10 cm-thick abdominal space, which was characterized using conductive signaling^17^ recorded from the subcutaneous space. We further observed increases of the plasma ghrelin levels, the “hunger hormone”, as the IngRI was activated *in vivo* to deliver a short-pulse stimulation (*A* ∼ 0.5 mA, *f* = 14 Hz, *pw* = 0.33 ms).

## RESULTS

### Contact and Stimulation

The multilayered IngRI (**Figure 2A**) consists of the following layers: (1) a bio-inspired adhesive layer made of lyophilized alginate-polyacrylamide (Alg-PAAm) hydrogel coated with ethyl(dimethylaminopropyl)carbodiimide (EDC)-N-hydroxysuccinimide (NHS)-chitosan bridge polymer (∼ 1000 µm in thickness) and soft silicone (Ecoflex 00-30, ∼ 100 µm in thickness) to achieve an adhesive interface with the gastric mucosa; (2) a stiff silicone-encapsulated functional layer (Sylgard 184 silicone, ∼ 500 µm in thickness) that supports a flexible, thin-film copper coil (∼ 18 µm in thickness) for receiving wirelessly transmitted power, which is then wire-bonded to a pair of stimulation electrodes; and (3) a small neodymium magnet (∼ 7.5 mm in diameter) to enable magnetic navigation after ingestion and to provide the necessary compressive forces for the onset of the hydrogel-enabled adhesive interface. A series of photolithography, transfer printing, and molding steps were used to fabricate the IngRI (see **Figure S1 and S2** and **Materials and Methods** for details). The resulting system is thin (∼ 1 mm in overall thickness excluding the magnet), soft (Et^3^ ∼ 1.71110^-3^ Nm, where E is the effective elastic modulus and t is the effective thickness), and stretchable (∼ 20% in uniaxial stretchability), and consists of discrete functional parts that fully address the C-P-M-R issues and can be folded and loaded into a 000-size commercial gelatin capsule to allow oral administration (**Figure 2B**).

**Figure 2:**
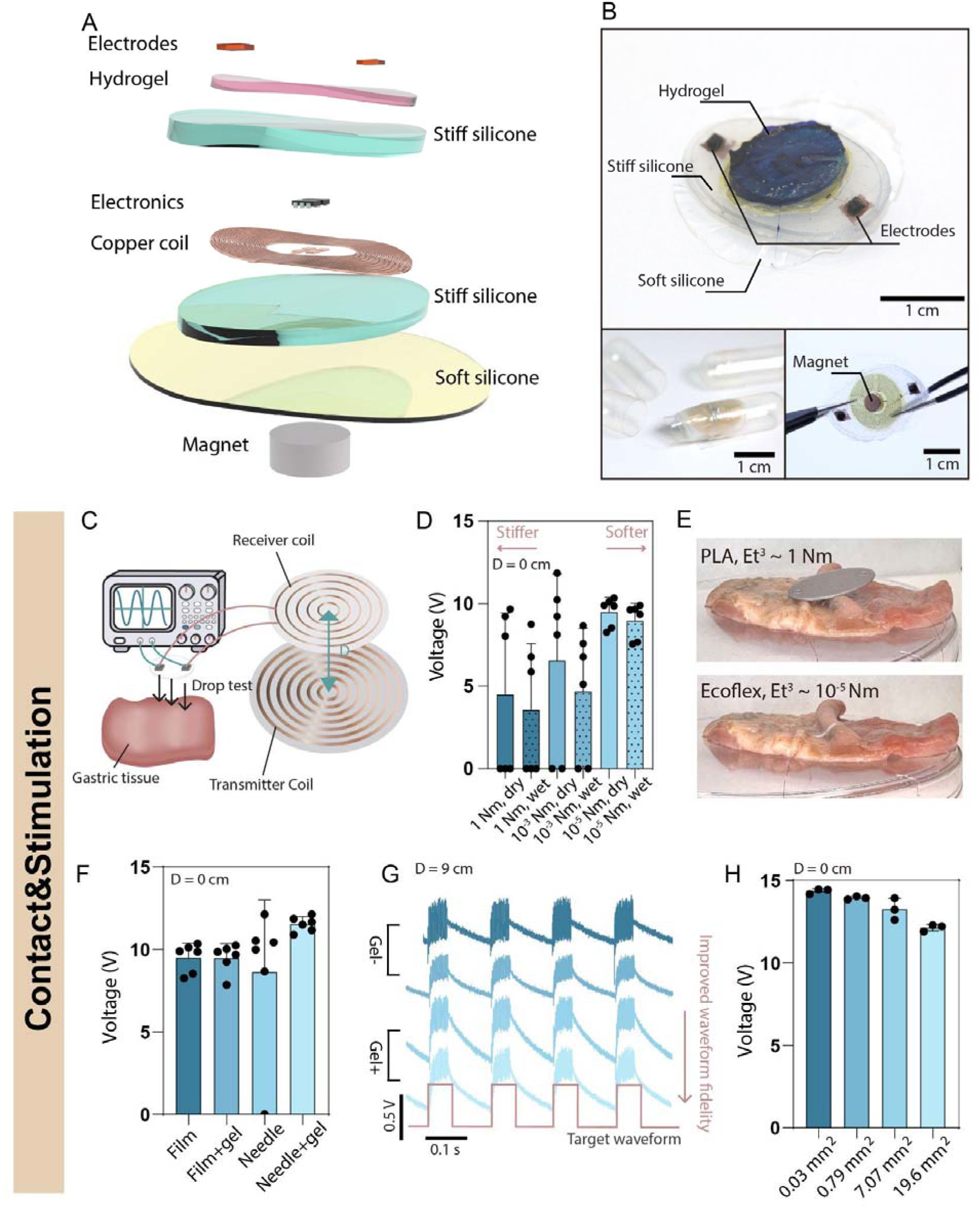
Design, contact, and stimulation properties of the IngRI. (A) Exploded, multilayered view of the IngRI. (B) Top: optical images of the IngRI with major components labeled. Bottom left: the IngRI folded and fitted inside a gelatin capsule. Bottom right: the IngRI deformed to a significant degree, demonstrating its flexibility. Scale bars, 1 cm. (C) Schematic showing the drop test setup: the IngRI is wired to the receiver coil which receives wireless power from the transmitter coil, and simultaneously connects to an oscilloscope to display output voltages under different controlled parameters. (D) Average output voltages of IngRIs with different stiffnesses as they are dropped onto wet or dry porcine gastric tissues. The result clearly shows that a smaller stiffness leads to both a higher average output voltage and a smaller error bar. (E) Optical images of the rigid (top) and soft (bottom) versions of the IngRI interfaced with the gastric tissue after dropping. (F) Average output voltages using different electrode materials and configurations. (G) Representative output pulses (blue color pulses) with (gel+) and without (gel-) PEDOT: PSS as the contact interface, compared with ideal waveform (brown color pulses). Electrodes with the gel show better waveform fidelity. (H) Average output voltage with different electrode areas, showing that the output voltage inversely scales with the contact area.

Through an *ex vivo* electrode drop test (**Figure 2C**), we showed that a soft device architecture with low flexural rigidity was better than a rigid one (which is the case for most existing IDEs) for establishing a high-fidelity electrode-tissue contact. This is particularly the case for gastric mucosa, which is approximately 100 times softer than skin^28^, has more folds and wrinkles, and experiences constant peristaltic motion. We attached the IngRI (receiver coil diameter = 22 mm, **Figure S3**) onto substrates with different rigidities (Et^3^ ∼ 10^-5^, 10^-3^, and 1 Nm, respectively) and dropped it in multiple (n>5) random locations of a segment of *ex vivo* porcine gastric tissue (dry or wet) to simulate actual *in vivo* deployment. At a constant transmission distance (D = 0 cm), the output voltage increased with less variability as the substrate became softer, with or without the fluid (**Figure 2D**). Visualization of the device-tissue interface confirmed this trend (**Figure 2E**), in which the stiffer IngRI frequently failed to achieve stable contact due to a mechanical mismatch with the underlying tissue.

The materials and configurations of the electrodes also influence the quality of the electrode-tissue contact. Four different types of electrodes were fabricated and tested on segments of *ex vivo* porcine gastric tissue: film (Cu, ∼ 6 µm in thickness), film + gel (poly(3,4-ethylenedioxythiophene) polystyrene sulfonate, or PEDOT:PSS, interpenetrated with polyacrylic acid ∼ 1 mm in thickness, see **Figure S4** and **Materials and Methods** for details), needle (stainless steel, ∼ 0.5 mm in diameter), and needle + gel (**Figure S5**). The results showed that the addition of PEDOT:PSS gel improved the amplitude and reduced the variation for both the film and wire electrodes (**Figure 2F**), and helped preserve the output waveform fidelity at long transmission distances (D = 9 cm, **Figure 2G**). This is likely due to a combination of mechanical (low modulus and stretchability) and electrical (dual ionic-electronic conductivity) properties of the PEDOT:PSS gel, which helps to lower the overall contact impedance^29,30^. Furthermore, the output voltage was found to be inversely proportional to the electrode area (**Figure 2H**).

### Navigation and Retention

To support navigation of an ingestible device we implemented a simple, low-cost, and non-invasive approach by attaching a small neodymium magnet to the IngRI, which allowed remote manipulation using a larger, hand-held neodymium magnet from outside the body. Using the *in vitro* setup shown in **Figure 3A**, we found that the minimum actuation distance while using a hand-held neodymium magnet (5 cm in diameter, 2.5 cm in thickness, ∼ 0.47 T in surface magnetic strength), scaled inversely with weight of the IngRI, and increased significantly if the IngRI was submerged in water (**Figure 3B**). Because the average depth from the gut to the skin surface of the upper abdomen is ∼ 2 cm in adult humans^31^ and ∼ 6 cm in adult Yorkshire pigs, this setup was more than adequate to test the navigation abilities of the IngRI in adult subjects. Indeed, in anesthetized swine models (see **Materials and Methods** for details), the IngRI can be moved across the majority of the gut to designated gastric positions using a large, handheld magnet placed against the chest of the swine (**Figure 3C and Movie 1**). Notably, the IngRI can be flipped inside the gut by reversing the polarity of the handheld magnet (**Figure 3D and Movie 2**), which is a critical feature for ensuring that the electrodes can always correctly face and contact the gastric mucosa.

**Figure 3:**
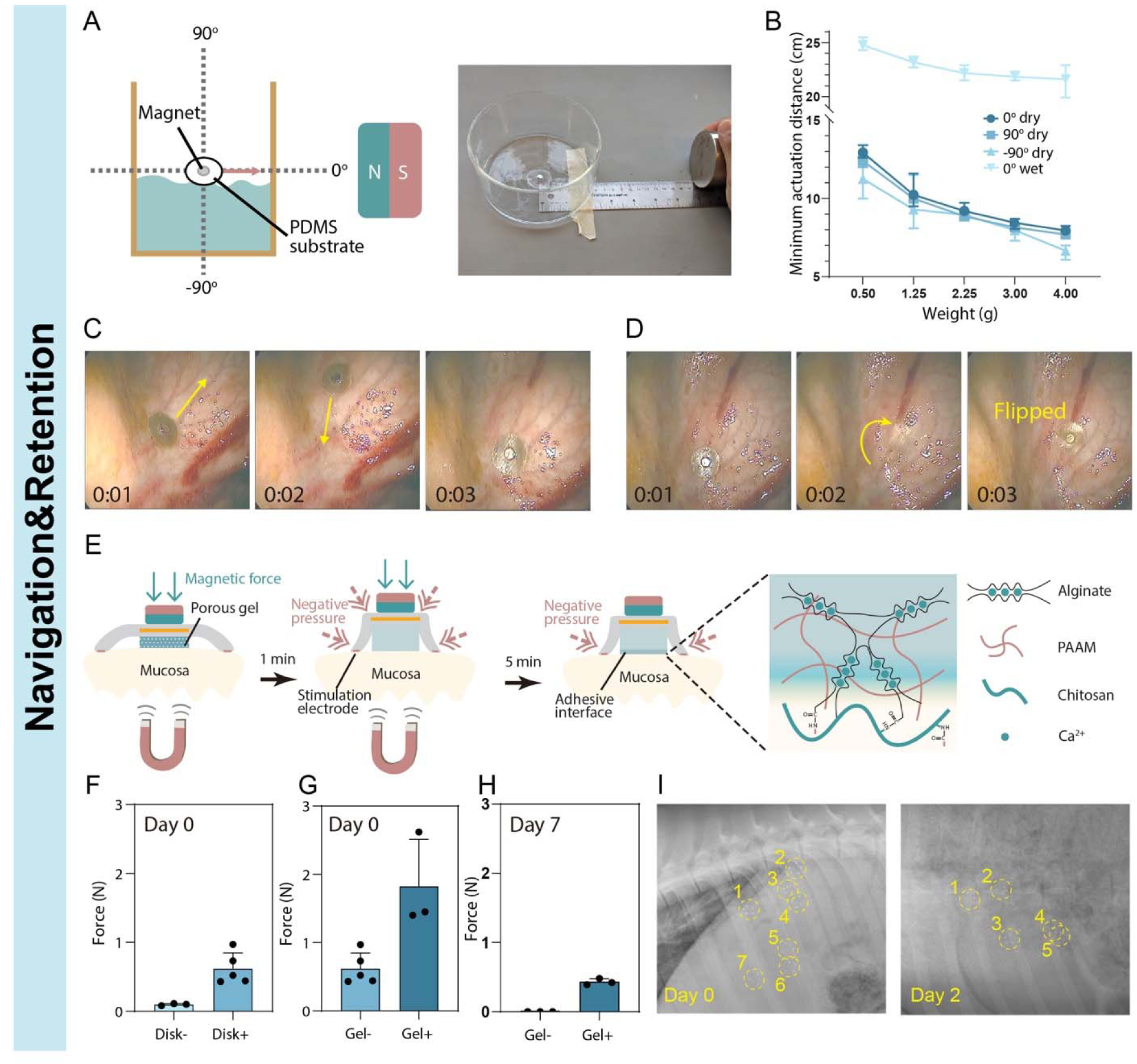
Navigation and retention properties of the IngRI. (A) Schematic (left) and optical image (right) of the in vitro setup for simulating in vivo navigation of the IngRI using an external hand-held neodymium magnet. (B) The minimum actuation distance of the IngRI when manipulated by a hand-held neodymium magnet under different orientations and wetness conditions obtained from setup in (A). (C-D) Demonstrations of in vivo navigating and flipping ability of the IngRI. Pictures are captured from Supplementary Movies 1 and 2. (E) Schematics of the bio-inspired hydrogel adhesive for enhancing gastric retention of the IngRI. Briefly, the mechanism relies on a bilayer of soft silicone disk and lyophilized hydrogel, which together provide a combination of physical and chemical adhesion to overcome the challenging gastric environment. (F) Average peeling forces of the IngRI with (disk+) and without (disk-) the soft silicone disk at day 0. (G) Average peeling forces of the IngRI with (gel+) and without (gel-) lyophilized hydrogel at day 0. (H) Average peeling forces of IngRI with (gel+) and without (gel-) the bio-inspired hydrogel adhesive at day 7. (I) Radiographs at day 0 and day 2, respectively, tracking the gastric retention of devices with the bio-inspired hydrogel adhesive in the porcine stomach. 5 out of 7 devices are intact after 48 hours of ingestion.

Apart from navigation, the ability to remain closely attached to a fixed gastric location for more than 48 h is crucial for realizing long-term gastric electrostimulation. However, this feature is not addressed by existing IDEs that have an average gastric retention time of less than 4 hours^32^. Inspired by leeches, we introduced a tissue-adhering interface in the form of a suction disk coated with a water-absorbing tissue-adhering lyophilized hydrogel adhesive. Leeches achieve strong adhesion to biological surfaces by creating negative pressure using disk-like muscle tissues^33^. Here, we recreated a similar disk-like soft structure, in which negative pressure was provided by the liquid absorption of the underlying lyophilized hydrogel. As depicted in **Figure 3E**, as soon as the structure contacts the wet mucosal surface, the lyophilized hydrogel begins absorbing liquid, whereas the surrounding suction disk blocks further liquid infusion. This immediately creates a negative pressure that results in physical adhesion between the disk and mucosa. After approximately 5-10 minutes, the bridge polymer began to interpenetrate the gastric mucosa to form covalent bonds between the primary amino and carboxyl groups^34^, resulting in strong chemical adhesion (**Figure S6**). This combination of physical and chemical bonding is key to realizing extended gastric retention and achieves superior results compared with using only one of these mechanisms alone.

Peeling tests were performed using *ex vivo* porcine gastric tissues, and the presence of disks and gels was found to increase the peeling force by 6-fold (**Figure 3F**) and 3-fold (**Figure 3G**), respectively, compared to the controls. The long-term stability of the tissue-adhering interface was evaluated by immersing IngRI attached to *ex vivo* porcine gastric tissues in phosphate-buffered saline. After 7 days, the resulting peeling force was reduced compared to that on day 0, but was still significantly larger than that of the control (**Figure 3H**). We noticed that the presence of the disk significantly increased the chemical bonding activity between the gel and mucosa because it provided physical and mechanical stability during the initial 5-10 minutes necessary for the onset of chemical bond formation, and 2) it created a sealing mechanism around the gel to further minimize water ingression and gel swelling. This was evident through visual examination at the tissue-gel interface after 7 days of soaking (**Figure S7**), where the gels with disks were ∼ 40±9% less swollen than those without disks. Furthermore, we evaluated long-term gastric retention of IngRI using ambulating swine models (see **Materials and Methods** for details): five out of seven devices delivered into the porcine gut remained intact while the swine were allowed to feed three times a day for 48 h, as shown by the radiographs (**Figure 3I**).

### Powering and Modulation

Existing IDEs are mostly powered by coin cell batteries and carry sophisticated on-board microcontrollers and wireless communication modules for the modulation of electrostimulation waveforms, which occupy a significant device volume, reduce device flexibility, and impose safety risks^14,35^. In this study, we developed a wireless, battery-free, and chip-free strategy for powering and modulating the IngRI inside the gut, which consisted of pulse wave inputs, high-power radio frequency (RF) transmission waves, and on-board receiving and rectifying circuits (**Figure 4A**). An Arduino Nano board was utilized to generate pulse wave signals with a fixed *f* = 14 Hz and adjustable pulse waves (PW) (**Figure 4B**). The PW can be tuned from 0.33 to 50 ms, which covers most reported waveforms used experimentally and clinically for electrostimulation of major organs (**Figure 1A**). The pulse wave signals were sent to a waveform generator (Keithley 3390, USA) to produce an 11 MHz RF signal using the frequency-shift keying (FSK) modulation scheme (**Figure 4B**), which was then amplified by an RF linear power amplifier (Bell Electronics, 2200 L) and loaded onto a portable transmitter coil (**Figure S8**). In the IngRI receiver coil, capacitor C_1_ was used for resonant frequency matching with the transmitter coil and capacitor C_2_ was used to stabilize the output voltage on the load. Two diodes, D_1_ and D_2_ acted as rectifiers to generate the unilateral pulsed waveforms (**Figure S9**). Three example output waveforms with fixed *f* = 14 Hz and different PWs recorded from R_load_ are shown (**Figure 4C**). The peak voltage for short PWs (e.g., 0.33 ms) dropped ∼ 5-fold compared to that for longer PWs (**Figure 4C**), likely due to a premature signal cut-off caused by the charging time of capacitor C_2_.

**Figure 4:**
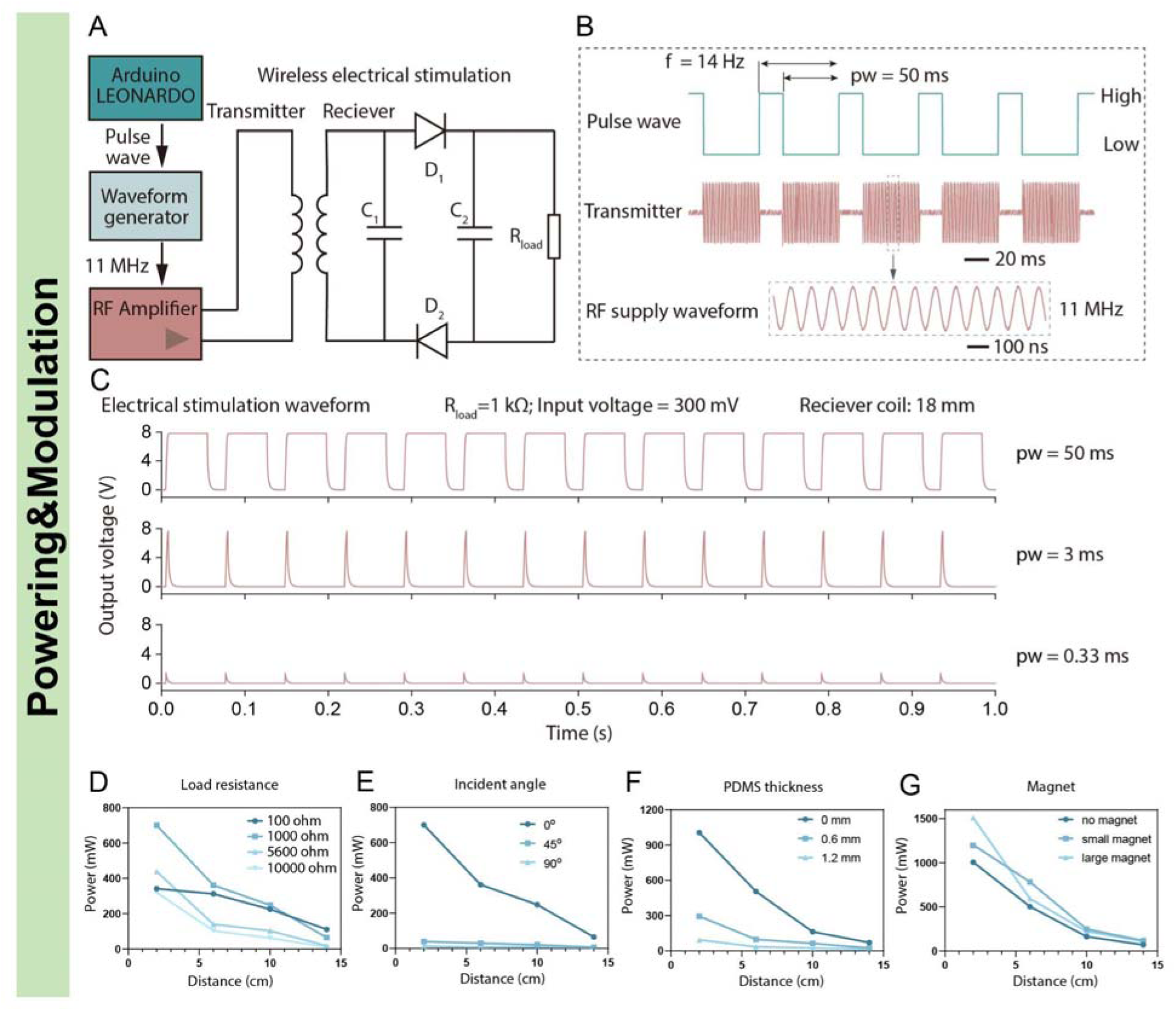
Powering and modulation properties of the IngRI. (A) Schematic showing the electronic setups of the near-field inductive coupling. (B) Conversion of a pulse wave into a high-frequency pulse train for wireless transmission to the receiver coil. (C) Representative output waveforms with frequency = 14 Hz, load resistance = 1000 ohm, and receiver coil diameter = 18 mm under different pulse widths (0.33, 3, and 50 ms). (D-G) The output power as a function of distance between the transmitter and receiver coils, under different (D) load resistances, (E) incident angles, (F) PDMS encapsulation thickness, and (G) presence of magnets.

Receiver coils with different diameters (14, 18, and 22 mm) were fabricated to investigate the effect of coil size on the output signals. Different capacitors (33, 8, and 1.5 pF) were used to match the resonant frequencies near 11 MHz (**Figure S10**). The output voltage increased as the size of the receiver coil increased (**Figure S11A**), which was expected because the efficiency of RF energy collection typically scales with the antenna size. The resonant frequency drift under bending was measured, which remained below 1% for up to 0.1 mm^-1^ bending curvature (**Figure S11B**), indicating good tolerance to mechanical deformations during *in vivo* deployment. The amplitude of the output voltage scaled linearly with the waveform generator’s input voltage for both PW = 50 ms and 0.33 ms (**Figure S11C**), demonstrating fine adjustability of the electrostimulation intensity using the external circuit.

Next, the efficiency of wireless power transfer subject to various engineering parameters was evaluated. Using an *in vitro* setup (**Figure S12**), the output power as a function of the distance between the transmitter and receiver coils was measured (**Figure 4D**), and a load resistance of 1000 Ω was found to produce the highest power output compared to other loads (**Figure 4D**), which is similar to the electrical impedance between the stimulation electrodes and *in vivo* gastric tissues^17^. The output power as a function of the incident angle between the coils was also measured (**Figure 4E**), and the optimal orientation was found to be parallel (i.e., angle = 0°). Finally, PDMS encapsulation was found to greatly attenuate the output power, whereas the presence of the central magnet slightly enhanced it (**Figure 4F&G**). Therefore, a PDMS thickness of ∼ 500 µm was selected in the final design of the IngRI to mitigate power loss while serving as a physical barrier against the acidic and aqueous gastric environments for at least a month.

### In vivo Evaluation

The efficacy of IngRI was evaluated in anesthetized swine models (**Figure 5A**). During the procedure, the IngRI was folded, loaded into a 000-size commercial gelatin capsule, and delivered through an overtube via the oral route into the gut, assisted by an endoscope. The capsule was dissolved within minutes to release the device, allowing magnetic navigation and positioning of the IngRI at the desired gastric location. The transmitter coil was then placed on top of the IngRI at an approximately parallel angle. In the first demonstration, a commercial light-emitting diode (LED, minimum operation power ∼ 70 mW) was connected to the IngRI to validate the wireless power transfer through the abdominal tissue. As shown in **Figure 5B**, a small input voltage from the waveform generator (sinusoidal wave, 300 mV p-p, *f* = 14 Hz, PW *=* 50 ms) was sufficient to light the LED inside the gut, which remained operational for at least an hour in the presence of food particles and gastric juice. The distances between the two coils varied with different swine sizes, and were estimated to be between 5 and 10 cm. As swine tissues are significantly thicker than those of humans, a considerably lower input voltage is required to operate the same LED inside the human gut.

**Figure 5:**
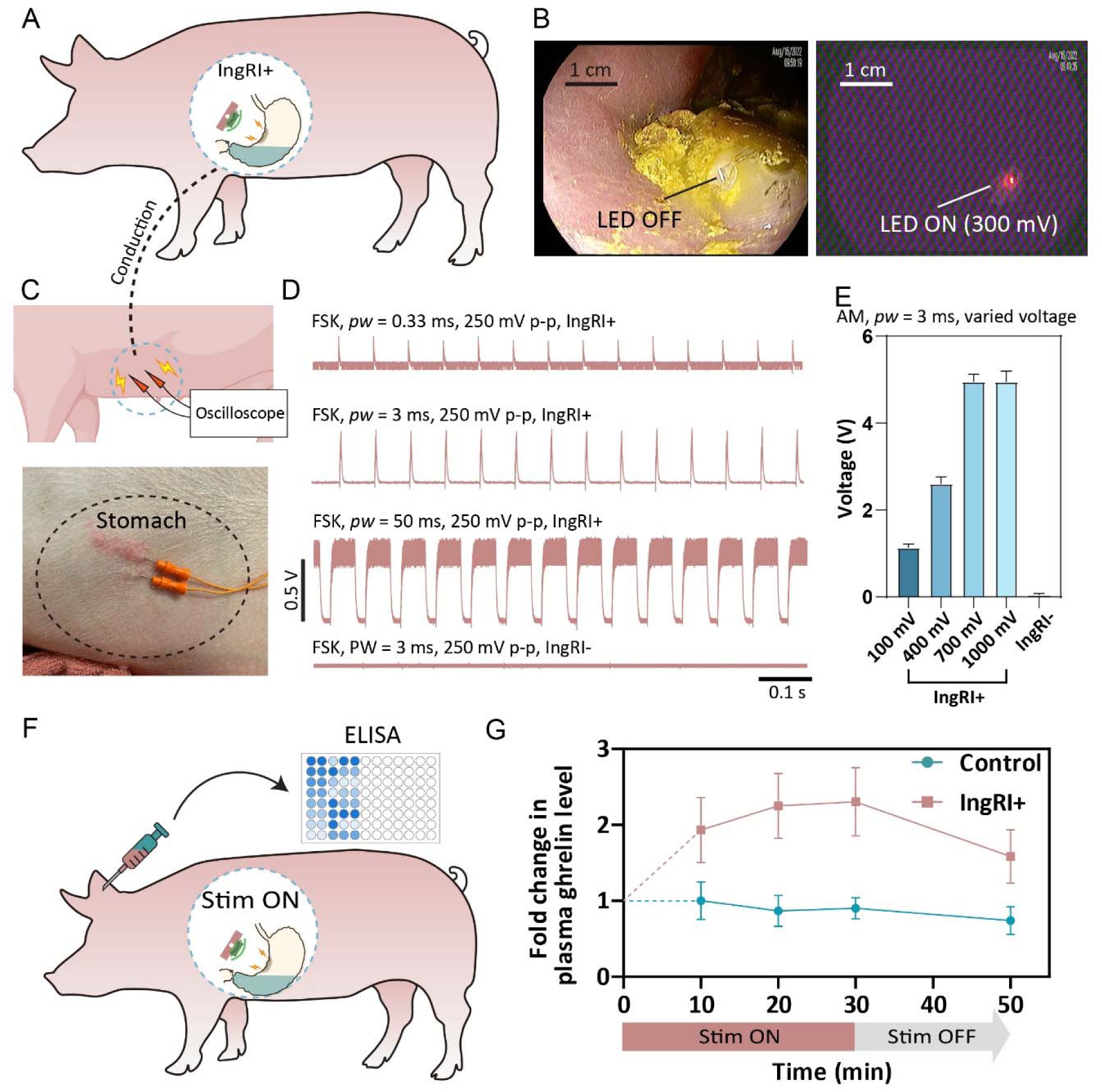
In vivo validations of the IngRI. (A) Schematic of in vivo experiments using anesthetized swine models. (B) Endoscopic images showing wireless powering of an LED device using the IngRI inside the gut. (C) Schematics of in vivo experiments to measure the conductive signaling delivered using the IngRI. Needle electrodes are placed at the swine’s abdomen to detect and record the corresponding waveform. (D) Recorded waveform from the subcutaneous needle electrodes as waveforms with different pulse widths are delivered under an FSK modulation scheme. A control experiment without the IngRI inside the gut is shown in the bottom to show that the wireless transmission did not interfere with the subcutaneous recording. (E) Output voltages of the subcutaneous recording as different input voltages are used, under an AM modulation scheme. (F). Schematic of plasma ghrelin level measurements in anesthetized swines stimulated with the IngRI. (G). Averaged (n = 3) fold change in plasma ghrelin levels as a function of time (IngRI, red; control, green).

Next, to validate the ability of IngRI to effectively deliver electrical pulses to the gastric wall, conductive signaling from the nearby subcutaneous space was recorded by inserting a separate pair of needle electrodes into the swine abdomen (**Figure 5C**). Representative waveforms were detected subcutaneously as the IngRI was turned on and tuned to different PWs (0.33, 3, and 50 ms) using the FSK modulation scheme (**Figure 5D**), which closely resembled the *in vitro* results in **Figure 3C**; for example, a reduced amplitude at a smaller PW (e.g., 0.33 ms) was similarly observed. A control experiment was also performed, in which the input voltage was turned on without the IngRI inside the gut, and no conductive signaling was observed (**Figure 5D**). By switching from an FSK modulation scheme to an amplitude modulation (AM) scheme, a much higher voltage output (∼ 3 times higher) was observed using the same input voltage, which became saturated at an input voltage of ∼ 700 mV p-p (**Figure 5E**). The maximum output voltage (∼ 5 V) translates to ∼ 5 mA of current if a tissue impedance of 1000 Ω is assumed, which is comparable to the clinically available, surgically implanted stimulators^34^.

To further evaluate the clinical potential of IngRI, plasma ghrelin levels were monitored for a 40-min period with the IngRI turned on for 30 min in a total of six swine models (**Figure 5F**, see **Materials and Methods** for details). During each experiment, blood was collected and analyzed simultaneously from two separate swine of similar weights and fed states (24 h of liquid diet), where one served as a control without IngRI. Under a short-pulse stimulation condition (*f* = 14 Hz, PW = 0.33 ms) and an estimated amplitude ∼ 0.5 mA, the plasma ghrelin levels of the swine with the IngRI were found to increase by ∼ 2-fold, from 93.9±23.4 pg/mL to 181.4±40.3 pg/mL, after 10 min of electrostimulation (**Figure 5G**). Plasma ghrelin levels continued to increase for the first 30 min and rapidly dissipated after the IngRI was turned off (**Figure 5G**). These findings are consistent with the literature results^36,37^, suggesting the potential of IngRI to regulate appetite and treat other endocrine conditions.

We further evaluated the safe passage of the device by feeding IngRIs without an adhesive interface to ambulating swine. Radiographs showed that they safely exited the GI tract within 2 days in 6 of the 6 trials (**Figure S13**), during which time the swine experienced no change in their dietary or sleeping patterns.

## DISCUSSION

This study demonstrates a proof-of-concept soft robotic interface (IngRI) for realizing ingestible, wireless, and acute electrostimulation of the GI tract. Both *ex vivo* and *in vivo* testing showed that a soft device architecture inspired by skin-interfacing wearable electronics yielded better electrode-tissue contact and longer gastric retention than the existing rigid IDEs. Short-pulse stimulation delivered by IngRI further resulted in an increase in plasma ghrelin levels, a gut hormone known to activate the growth hormone secretagogue receptor (GHS-R) and stimulate food intake and fat deposition^38^. This data support the potential for regulating appetite and treating other conditions through non-pharmaceutical neuromodulation, which could potentially result in a reduced side effect profile^39^. Future studies in large mammals and humans are needed to evaluate the optimized parameters of electrostimulation and identify the therapeutic window for treatment.

A notable potential advantage of IngRI is its capability for chronic (up to a few days) gastric stimulation while being completely ingestible. This was achieved through a combination of optimized wireless energy-transfer schemes and bio-inspired adhesive technologies that can be remotely triggered and interrogated. The IngRI can be compared to recently published, state-of-the-art ingestible electronics for providing electrical or mechanical stimulations, such as RoboCap^40^, STIMS^17^, and FLASH^41^, which have similar power consumption on the order of 100 mW, but are battery-powered and drain quickly *in vivo* (between 15 to 30 min), and cannot provide gastric retention for more than 24 h in ambulating large animal models. The extended *in vivo* lifetime of IngRI can significantly enhance efficacy by enabling modes of chronic modulation that are more akin to implanted gastric stimulators, which is the current clinical standard^42^, while improving patient acceptance and reducing costs. The long-term correlation between electrostimulation and physiological changes, such as weight loss and dietary changes, should be evaluated in more comprehensive clinical trials using large mammals and humans.

Similar to a wearable electronic platform, the IngRI can serve as a device chassis to enable a wide range of medical applications targeting the GI mucosal surface. For instance, stimulation electrodes can be exchanged for miniaturized vibrational, optical, or thermal sources to enhance drug absorption through the mucosa^40^ or to enable optogenetic induction of GI motility^43^. IngRI may also integrate miniaturized sensors to allow closed-loop and on-demand triggering of stimulators based on real-time physiological feedback, such as during micromanagement of dietary cycles in obese patients.

Future iterations of device design should address several specific safety, environmental, and technological concerns to assist clinical translation in humans. First, the presence of small neodymium magnets can interfere with imaging techniques involving magnetic fields (e.g., magnetic resonance imaging). However, several commercialized video capsule endoscopies introduced by Stereotaxis^44^ have overcome this limitation, and such strategies may be employed here. Second, although most materials of IngRI (silicone, copper, epoxy) are FDA-approved for ingestible devices, they are not biodegradable and therefore pose potential environmental concerns when excreted. Methods to retrieve and properly dispose of IngRI from excreted waste need to be developed; alternatively, they can be made using biodegradable electronic materials with designated degradation times to avoid generating electronic waste. Finally, as wireless technology advances, the existing bulky and cabled wireless transmission system may be miniaturized and made portable to allow outdoor use of IngRI.

In conclusion, IngRI is a battery-free, magnetically-supported gastric-adherent, and orally administrable robotic interface capable of delivering multiple days of electrostimulation to the gut in swine models, leading to therapeutically meaningful changes in plasma ghrelin levels. We expect future iterations of the IngRI to enable portable use in out-of-clinic settings, extended lifetime to weeks or even months, and integration with sensors to realize closed-loop neuromodulation and treatment.

## MATERIALS AND METHODS

### Fabrications of the IngRI

#### Fabrication of the receiver coil

A patterned flexible copper coil on a polyimide (PI) film was fabricated as the receiver coil using photolithography. Briefly, a positive photoresist (AZ-5214, AZ Electronic Materials) was spin-coated on a thin copper-clad polyimide film (18 μm-thick Cu and 12.5 μm-thick PI) at 3000 rpm for 30 s and soft baked at 110 □ for 3min. Patterns of the receiver coil were obtained by ultraviolet (UV) exposure, followed by development in a metal-ion-free developer (AZ300 MIF, AZ Electronic Materials) and wet etching in FeCl_3_ solution. Individual devices were then cut from the large sheet using a laser processing machine (ProtoLaser U4, LPKF Laser & Electronics).

Following receiver coil patterning, capacitor C_1_ was soldered at both ends of the receiver coil to obtain a matching resonant frequency with the transmitter coil. Two diodes (D_1_ and D_2_) were soldered to act as rectifiers to limit the flow of the current in one direction. Then, capacitor C_2_ with a capacitance value of 2.2 μF was soldered to stabilize the output voltage in the high-frequency pulse width range. The solder joints were encapsulated using UV-curable epoxy (Loctite 4305).

#### Fabrications of the PEDOT:PSS electrodes

Interpenetrating network PEDOT:PSS hydrogels were synthesized as previously described^30^ using a secondary network monomer solution comprising acrylic acid, water, bisacrylamide, and a thermally activated radical initiator (200:800:2:1 wt ratio). A PEDOT:PSS gel was first formed by mixing the PEDOT:PSS solution with 10x phosphate buffered saline in a 9:1 vol:vol ratio and then poured into a cylindrical mold. Subsequently, the gel was immersed in an equivalent volume of acrylic acid monomer solution, which was exchanged with a fresh monomer solution three times over 24 h. Finally, the PEDOT:PSS gel encapsulating the monomer solution was placed in an oven at 70 °C oven and allowed to cure for 1 h. The film was cut into the desired dimensions from the resulting gel using a razor blade.

#### Fabrications of the tissue-adhering hydrogel interface

The Alg-PAAm tough hydrogel was prepared following a modified protocol based on a previous report^46^. Briefly, sodium alginate powder and acrylamide were dissolved in distilled water to a concentration of 2 wt. % and 12 wt. % respectively, and stirred overnight until a clean solution was obtained. After degassing, 10 mL of the precursor solution was mixed with 36 μL of a 2 wt. % N,N′-Methylenebisacrylamide (MBAA) and 8 μL of tetramethylethylenediamine (TEMED) in one syringe (BD, 20mL). 226 μL of 0.27 M ammonium persulfate (APS) and 191 μL of 0.75M CaSO_4_ slurries were injected into another syringe (BD, 20 mL). All the bubbles were removed before further cross-linking. After connecting by a female luer × female luer adapter (Cole-Parmer), solutions in two syringes were mixed by pushing the syringe pistons forward and back 10 times. The mixture was stored overnight in a closed glass mold at room temperature to allow complete polymerization. The cured hydrogel was bonded to a primer-activated suction disk using a glue. The entire system was stored in a −80□°C freezer overnight and then lyophilized for 12 h to freeze-dry the hydrogel. Before mucosal bonding, the dried hydrogel surface was treated with a mixture of bridge polymer and coupling reagents for the carbodiimide coupling reaction. The bridge polymer, chitosan, was dissolved in distilled water at 2.0 wt. % and the pH was adjusted to 5.5∼6 by acetic acid. EDC and NHS were used as coupling reagents. The final concentration of EDC and NHS in the bridging polymer solution was 12 mg/mL.

The outer lip structure of the suction disk was fabricated using the following procedure: First, poly(methyl methacrylate) (PMMA 950 K A4, MicroChem) was spin-coated on an 1-inch silicon wafer at 3,000 rpm and baked at 180 °C for 3 min. Then, a well-mixed Ecoflex 00-30 liquid precursor was spin-coated onto the wafer at 250 rpm and baked at 90 °C for 90 min. The resulting elastomeric film could be easily peeled off from the silicon wafer for subsequent assembly onto the suction disk.

#### Assembly of the IngRI

Copper electrodes were laser-cut from copper sheets (∼ 18 µm in thickness) and electrically connected to the receiver coil using 36-gauge wires. The solder joints were encapsulated using a UV-curable epoxy (Loctite 4305). After placing the coil into an oval mold, PDMS with a 10:1 mixing ratio was poured into the mold, with electrodes sticking out of the liquid surface. After curing the PDMS at 65 °C, copper electrodes were attached to the PDMS surface using a UV-curable epoxy (Loctite 5055). The outer lip structure was treated with a primer agent (Loctite SF 7701) and attached to the opposite side of the coil device by using a thin layer of super glue (Loctite Prism 4011). The same side was coated with hydrogel to complete the suction disk structure. An acrylic circle ring mold was temporarily attached to the bottom of the coil and the hydrogel precursors were poured onto the coil. The hydrogel was cured under UV exposure for 20 min. After detaching the mold, the IngRI was freeze-dried for several days to promote adhesion between the coil body and hydrogel film. An illustration of the assembly procedure is shown in **Figure S2**.

### *Ex vivo* evaluation of the IngRI

#### Experimental setup

The Arduino was connected to the AUX port of a waveform generator (Keithley 3390, USA). The modulated output of the waveform generator was then connected to the input of an RF amplifier (Bell Electronics, 2200 L). The output of the RF amplifier was connected to the transmitter coil, which was clamped onto a stand to maintain a precise relative angle and distance with the IngRI. The receiver coil was laminated onto a piece of freshly collected porcine stomach tissue with two electrodes facing down and in contact with the tissue. Two additional wires were used to electrically connect the two electrodes and ports of the oscilloscope. A detailed schematic of the setup is shown in **Figure 2C**.

#### Adhesion strength measurements

A manual mechanical testing stage (Mark-10, Model ES20) coupled with a force gauge (Mark-10, Model M4-05) was used to apply a precisely controlled tensile pulling force to the suction disks, which was then converted to adhesive strength using the contact area.

### Tissue and *in vivo* experiments

All animal experiments were conducted in accordance with the protocols approved by the Committee on Animal Care at the Massachusetts Institute of Technology. Swine models were chosen because of their anatomical similarity to humans in GI-related studies^47^. Randomization of the animals was not performed. Female Yorkshire swine (Cummings Veterinary School at Tufts University in Grafton, MA) weighing 30–70□kg and aged 3–6 months were used. The swine were placed on a liquid diet 24□h before the study and fasted on the day of the procedure. On the morning of the procedure, the swine were sedated using intramuscular injection of either 5□mg□kg^−1^ telazol (tiletamine/zolazepam), 2□mg□kg^−1^ xylazine, or 0.25□mg□kg^−1^ midazolam and 0.03□mg□kg^−1^ dexmedetomidine. After intubation, anesthesia was maintained with isoflurane (2–3% oxygen). Under anesthesia, vital signs were monitored and recorded every 15□min throughout the study. After the study, swine were woken up using intramuscular injection of 0.1□mg□kg^−1^ atipamezole and monitored closely during the recovery process until full recovery was achieved.

For studies on ambulating animals to monitor long-term gastric retention of the IngRI, the swine were fed normally and anesthetized on days 0, 3, and 5 for radiography.

For all *ex vivo* studies, gastric tissue collections were performed within 10 min of euthanasia, and the tissues were maintained in Krebs buffer and stored at 4 °C fridge during use.

### In vivo hormone study

To measure the effect of electrostimulation by IngRI on plasma ghrelin levels, we administered the device to anesthetized swine. Approximately 5 mL of blood was sampled every 10 min during a 40-min electrostimulation session from the intravascular ear catheter. Following the extraction, 300 μL of 4-(hydroxymercuri)benzoic acid sodium (10 mM in PBS) was added to the blood and immediately centrifuged at 4 °C for 10 min. The resulting plasma was mixed with 10% (by volume) of 1N hydrochloric acid and stored at −80 °C before enzyme-linked immunosorbent assay (ELISA).

The ELISA was performed using a ghrelin-acylated porcine ELISA kit (Biovendor, RA594062400R). After thawing the plasma samples from −80 °C to room temperature, they were diluted 1:1 in a dilution buffer provided by the kit. Next, samples in duplicate were dispensed into antibody-coated microtiter strips, and 100 μL of acylated ghrelin conjugate solution was added. The strips were covered with a cover sheet and incubated for 18 to 20 hours at 4°C. After incubation, the strips were washed with wash buffer three times, and 200 μL of Ellman’s reagent was added to each well. The plate was covered with an aluminum sheet and incubated in the dark at room temperature. The absorbance of each well was measured at wavelengths between 405 nm and 414 nm. The absorbance was measured every 30 min until the maximum absorbance reached a minimum of 0.5 A.U., blank subtracted. The average absorbance for each standard and sample well was calculated, and a standard curve (ghrelin concentration versus absorbance) was plotted. Plasma ghrelin levels were calculated based on the standard curves.

## Supporting information

Supplemental Material Combination

Attachment Video

Navigation Video

Release Video

## FUNDING

This work was supported in part by the Karl van Tassel (1925) Career Development Professorship and Department of Mechanical Engineering, MIT. K.N. acknowledges start-up funding for the ZJU100 professorship from Zhejiang University. B.Y. acknowledges fellowship from Banting Postdoctoral Fellowship, and NSERC Postdoctoral Fellowship. This work was also supported in part by the Foundation of National Natural Science Foundation of China (NSFC) (Grant No. 61421002) and City University of Hong Kong (Grant Nos. 9678274, 9667221, and 9680322).

## AUTHOR CONTRIBUTIONS

K.N., K.W., D.L., B.Y., X.Y., and G.T. conceived and designed the IngRI. K.N., K.W., D.L., B.Y., V.R.F., K.Y., J.Z., and J.L. fabricated the devices and performed the material, mechanical, and electronic characterization. K.N., J.C.M., and S.W. performed the in vivo experiments with the assistance of J.J., K.I., J.K., W.A., A.H., and K.R.. K.N., X.Y., and G.T. provided the funding and supervised the project. All authors contributed to the writing of the manuscript.

## COMPETING INTERESTS

There are no competing interests related to this work at the time of its submission. Complete details of all relationships for profit and not for profit for the G.T. can be found in the following link. https://www.dropbox.com/sh/szi7vnr4a2ajb56/AABs5N5i0q9AfT1IqIJAE-T5a?dl=0.

## DATA AND MATERIALS AVAILABILITY

All data associated with this study are presented in the manuscript or Supplementary Materials.

