## Supplemental Material Combination for "An ingestible, battery-free, tissue-adhering robotic interface for non-invasive and chronic electrostimulation of the gut"

**Table of contents – Supplementary Information**

Note S1. Soft base adhesion force induction
Note S2. RF decay in tissue

Figure S1. Schematic step-by-step fabrication procedure for a flexible, thin-film copper film.
Figure S2. Schematic step-by-step assembly procedure of IngRI.
Figure S3. Top view of the 2D receiver coil designs used in this work (diameters = 14,18, and 22 mm).
Figure S4. (A). Schematic step-by-step chemical synthesis of PEDOT:PSS. (B). Scanning electronic microscope images of resulting PEDOT:PSS cross-section.
Figure S5. Optical images of IngRI with different types of electrodes for drop test.
Figure S6. Optical image showing degrees of hydrogel swelling with or without the thin silicone disk after soaking in PBS for 7 days. Sample with disk (right) shows significantly less swelling.
Figure S7. Optical image of the transmitter coil.
Figure S8. The circuit design (left), and the optical image (right) of the receiver coil.
Figure S9. Resonant frequency characterizations for coils of different diameters (14, 18, and 22 mm) under different bending curvatures. The peak impedance indicates the resonant frequency for each condition.
Figure S10. Characterizations of transmission properties. (A) Output voltage of the coil with different diameters (14, 18, and 22 mm) under the same pulse input. (B) The resonant frequency of receiver coils with different diameters (14, 18, and 22 mm) at different bending curvatures. (C) Output voltage as a function of input voltage with different pulse width (0.33 and 50 ms) as input pulses.
Figure S11. Optical image of in vitro coil characterization setup. Distance between the transmitter and receiver coil is measured using a ruler placed perpendicular to the two coils.
Figure S12. Radiographs of the ingested IngRI in the porcine stomach without the adhesive interface. On day 2, all devices safely exit the GI tract.

Movie S1. IngRI showing ability of navigation and flipping.
Movie S2. Endoscopic inspection after attachment

**Supplementary Notes**

**Note S1**. Soft base adhesion force induction

Flexural rigidity describes the easiness for a plate to deform, or bent – the larger thickness, or large young’s modulus it is, the greater the rigidity, according to

$D:=\frac{H^{3}*E}{12(1-\nu^{3})} (1)$

where $H$ is the thickness, $\nu$ is the Poisson’s ratio, $E$ is the young’s modulus.

Referring to Laplace pressure, which is the difference between the inside and outside of a curved surface that forms boundary between two fluid regions. As the two media shares the same contacting surface, it can be further simplified to:

$$\Delta P=\gamma\left( \frac{1}{R1}-\frac{1}{R2} \right)=\gamma\left( 2*\frac{1}{R1} \right) (2)$$

where $R1$ and $R2$ are the radii of the contacting surfaces.

The curvature is proportional to Flexural rigidity and inversely proportional to Moment. The larger Flexural rigidity it is, the less Moment it gains at the same point, generally. As the plate is less rigid, the greater area it contacts the gastric surface, with higher pressure difference. Thus, soft interface guarantees large contacting forces due to Laplace pressure. according to

$M\left( x \right)= -D* \frac{d^{2}w}{dx^{2}}= -D*\frac{1}{R} \to\frac{1}{R}=-\frac{D}{M\left( X \right)} (3)$

where $M(x)$ is the moment at given point.

**Note S2**. RF decay in tissue

In the actual application of wireless power transfer based on near-field inductive coupling, received power is much smaller when the transfer medium is tissue, instead of air.

Notation:

μ_0=4π×10^(-7) H/m: vacuum permeability;

ϵ_0=8.854×10^(-12) F/m: vacuum permittivity;

ϵ_r: relative permittivity;

σ: conductivity (S/m);

ω: angular frequency (rad/s);

f: frequency (Hz);

According to Beer-Lambert law, the intensity I of an electromagnetic wave into a type of material drop exponentially with the distance from the surface z:

$$I\left( z \right)=I_{0}e^{-\alpha z} (1)$$

The expression of attenuation coefficient α is:

$$\alpha=\sqrt{\frac{\mu_{0}\epsilon_{0}\epsilon_{r}}{2}[\sqrt{1+\frac{\sigma^{2}}{\omega^{2}{(\epsilon_{0}\epsilon_{r})}^{2}}}-1]} (2)$$

Corresponding dielectric properties of body tissue at f = 13.56 MHz:

| Tissue type | Conductivity $\sigma$ (S/m) | Relative permittivity $\epsilon_{r}$ |
| --- | --- | --- |
| Fat | 0.03 | 11.827 |
| Muscle | 0.628 | 138.44 |

**Table S1**: Conductivity and relative permittivity of different tissues

Considering the scenario of transmitting wireless power at 13.56 MHz through the belly of a pig (from skin to GI tract), the medium is modelled as 3 cm-thick fat and 6 cm-thick muscle. Based on Eq. 1 and 2, the intensity of electromagnetic wave falls off to 70% of its original value at the surface. Since the power is the square of intensity, the transmitted power in tissue is 49% of that in air.

**Supplementary Figures**

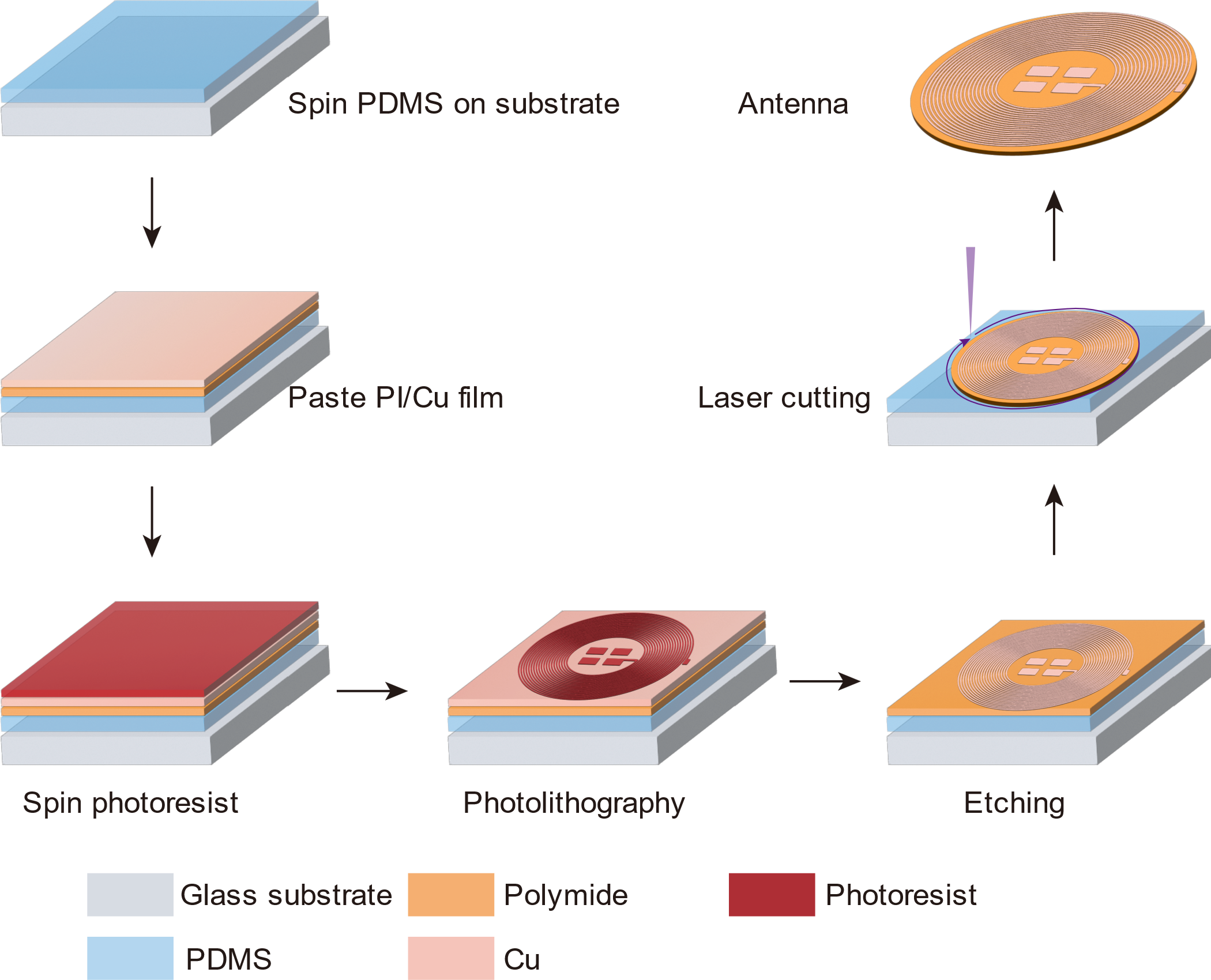

**Figure S1**. Schematic step-by-step fabrication procedure for a flexible, thin-film copper film.

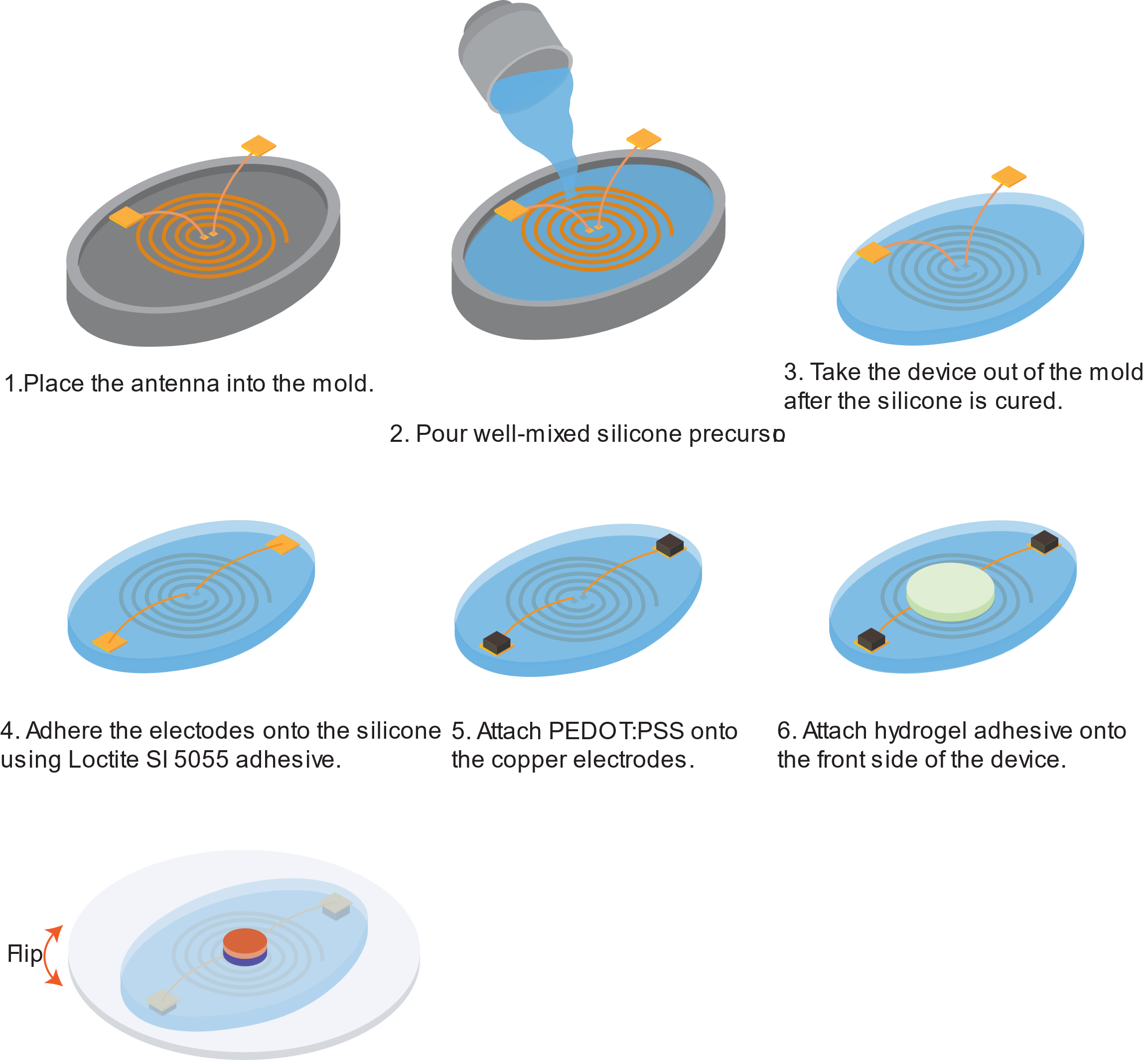

**Figure S2.** Schematic step-by-step assembly procedure of IngRI.

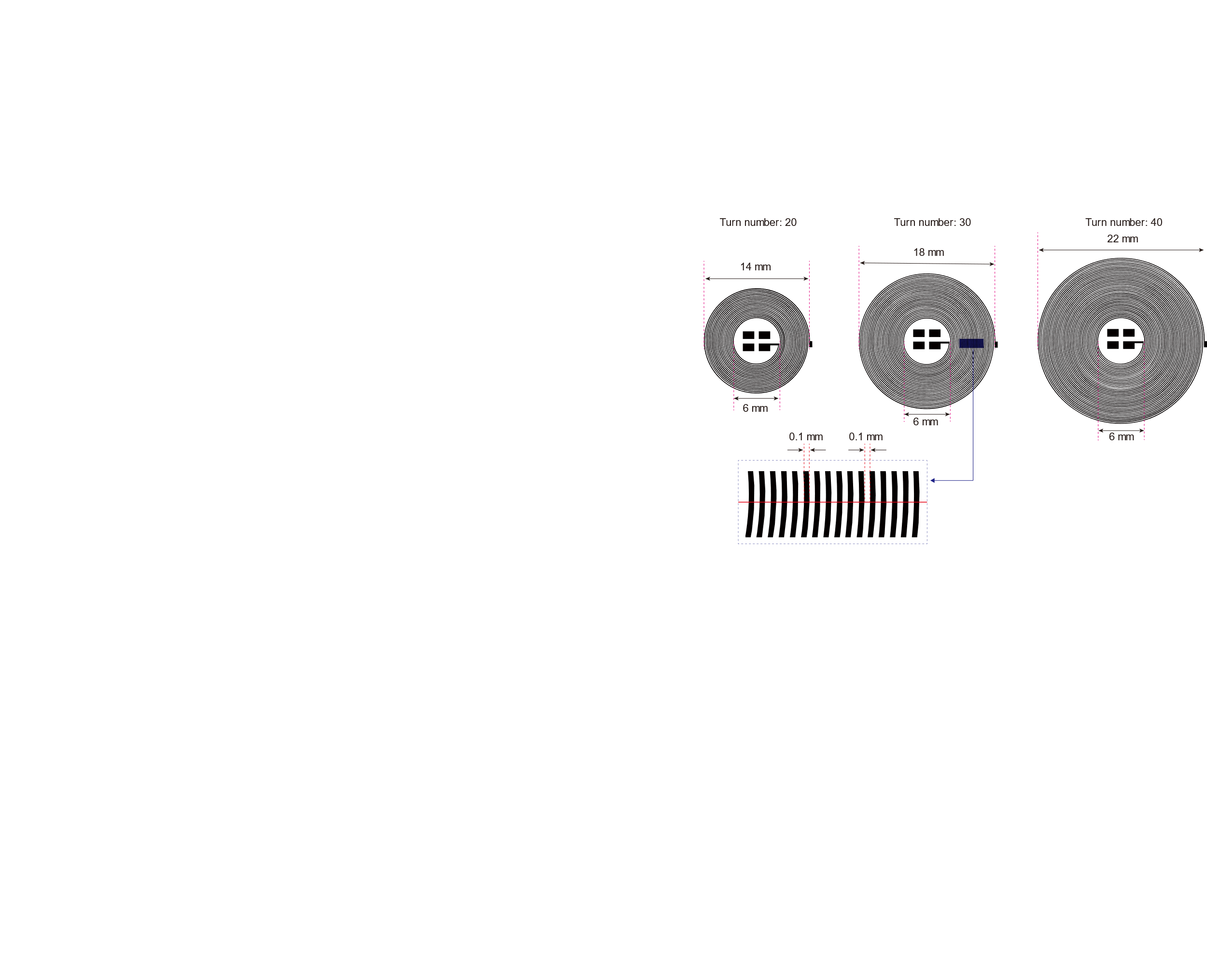

**Figure S3.** Top view of the 2D receiver coil designs used in this work (diameters = 14,18, and 22 mm).

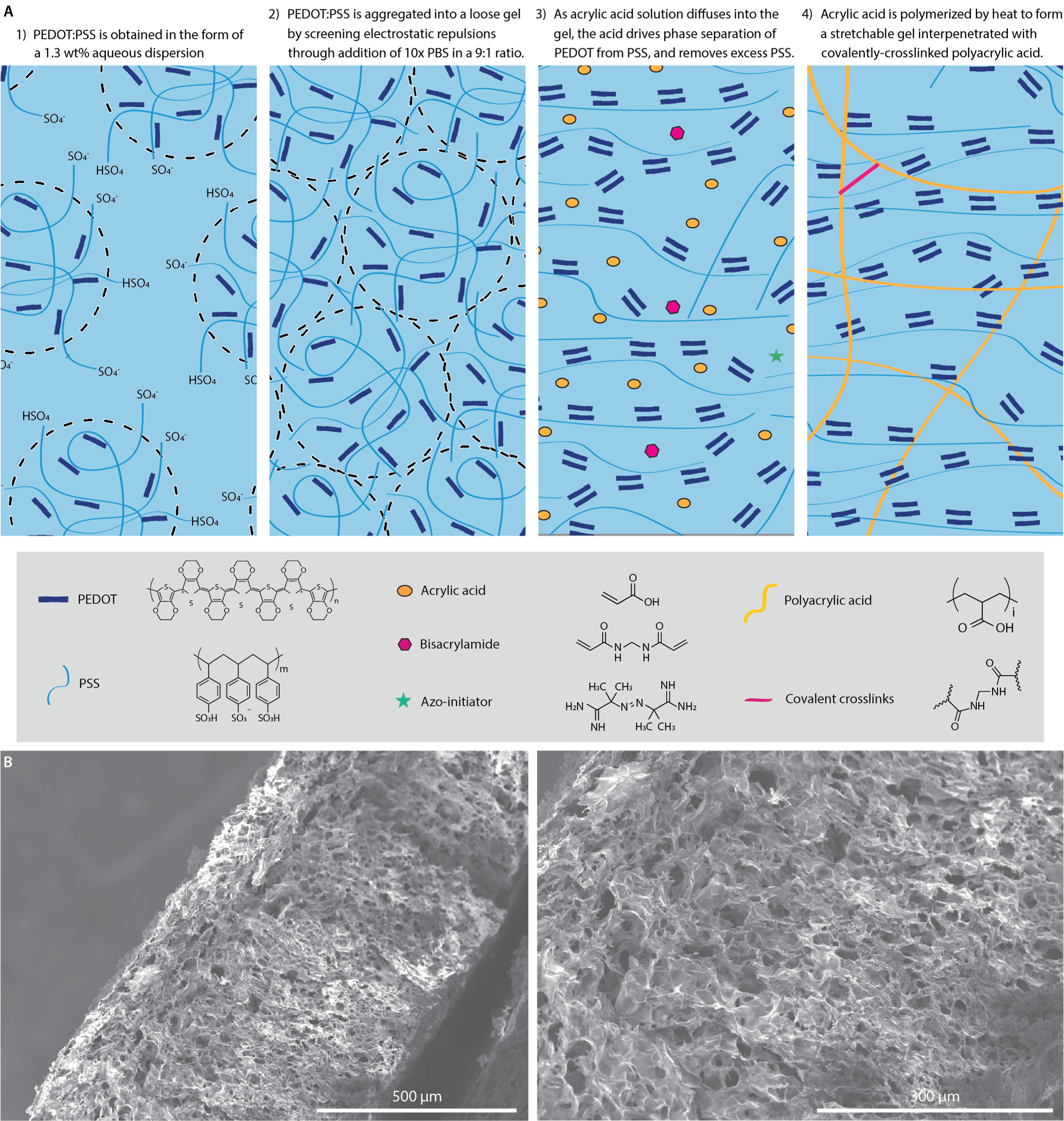

**Figure S4.** (A). Schematic step-by-step chemical synthesis of PEDOT:PSS. (B). Scanning electronic microscope images of resulting PEDOT:PSS cross-section.

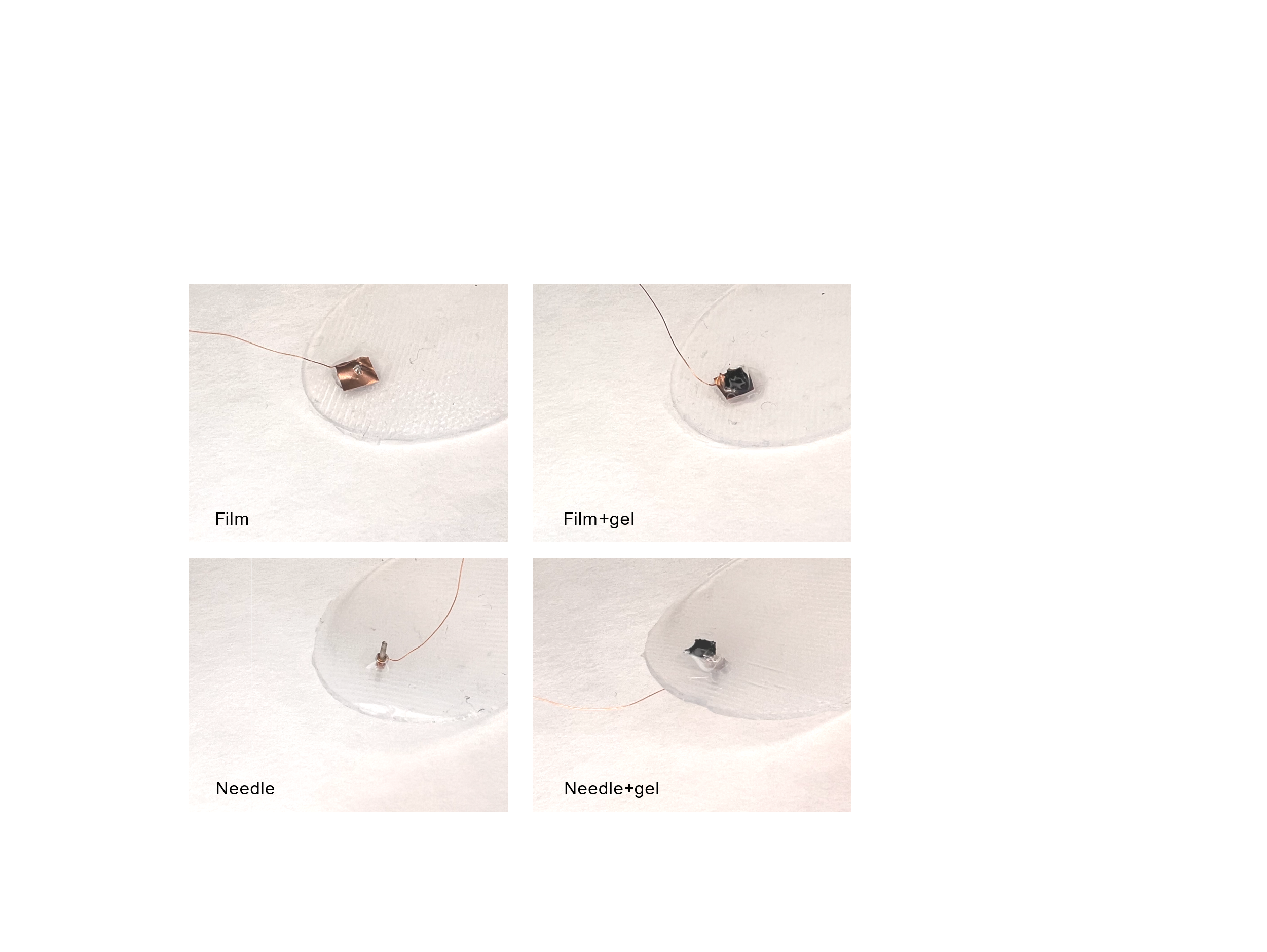

**Figure S5.** Optical images of IngRI with different types of electrodes for drop test.

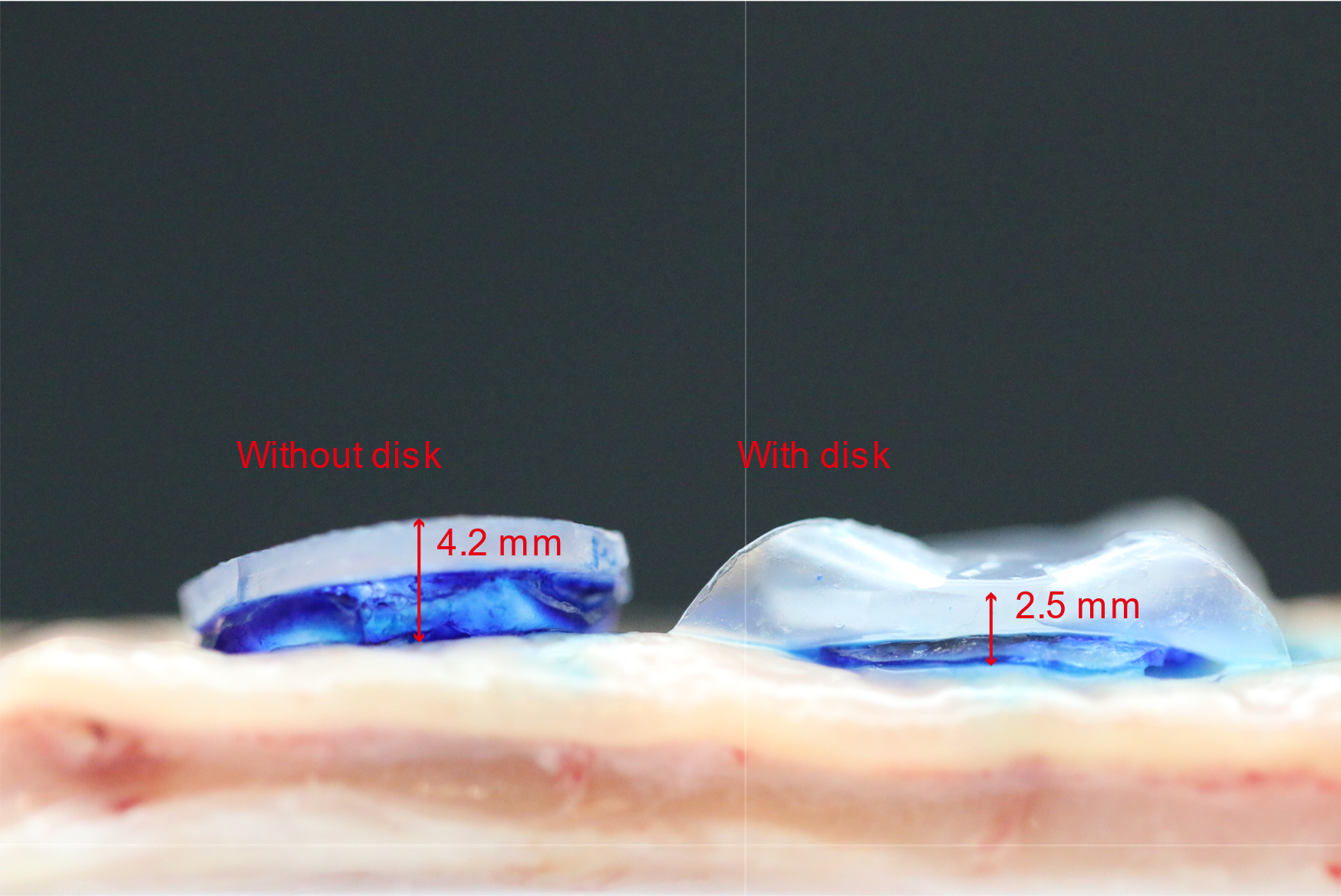

**Figure S6.** Optical image showing degrees of hydrogel swelling with or without the thin silicone disk after soaking in PBS for 7 days. Sample with disk (right) shows significantly less swelling.

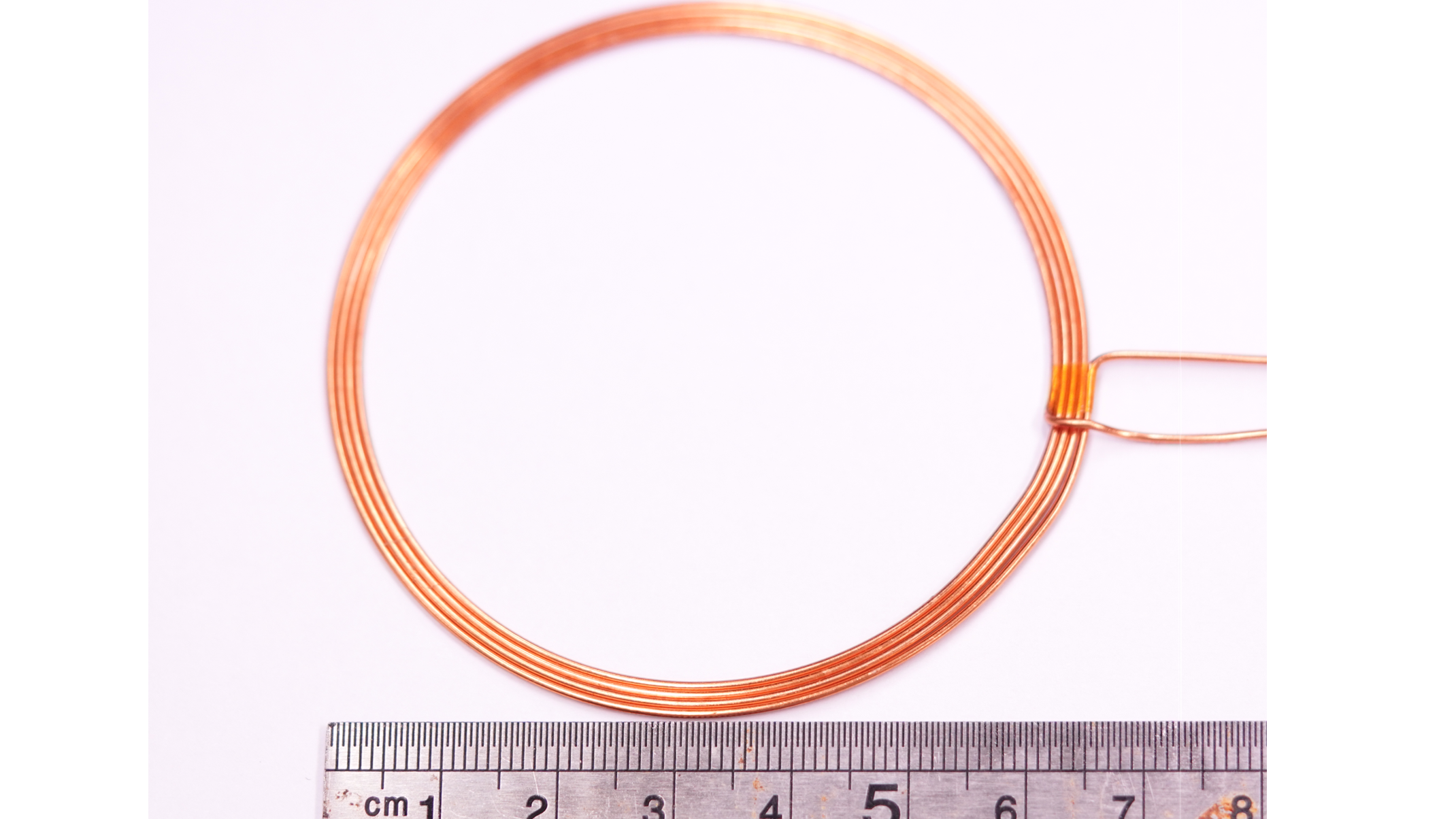

**Figure S7.** Optical image of the transmitter coil.

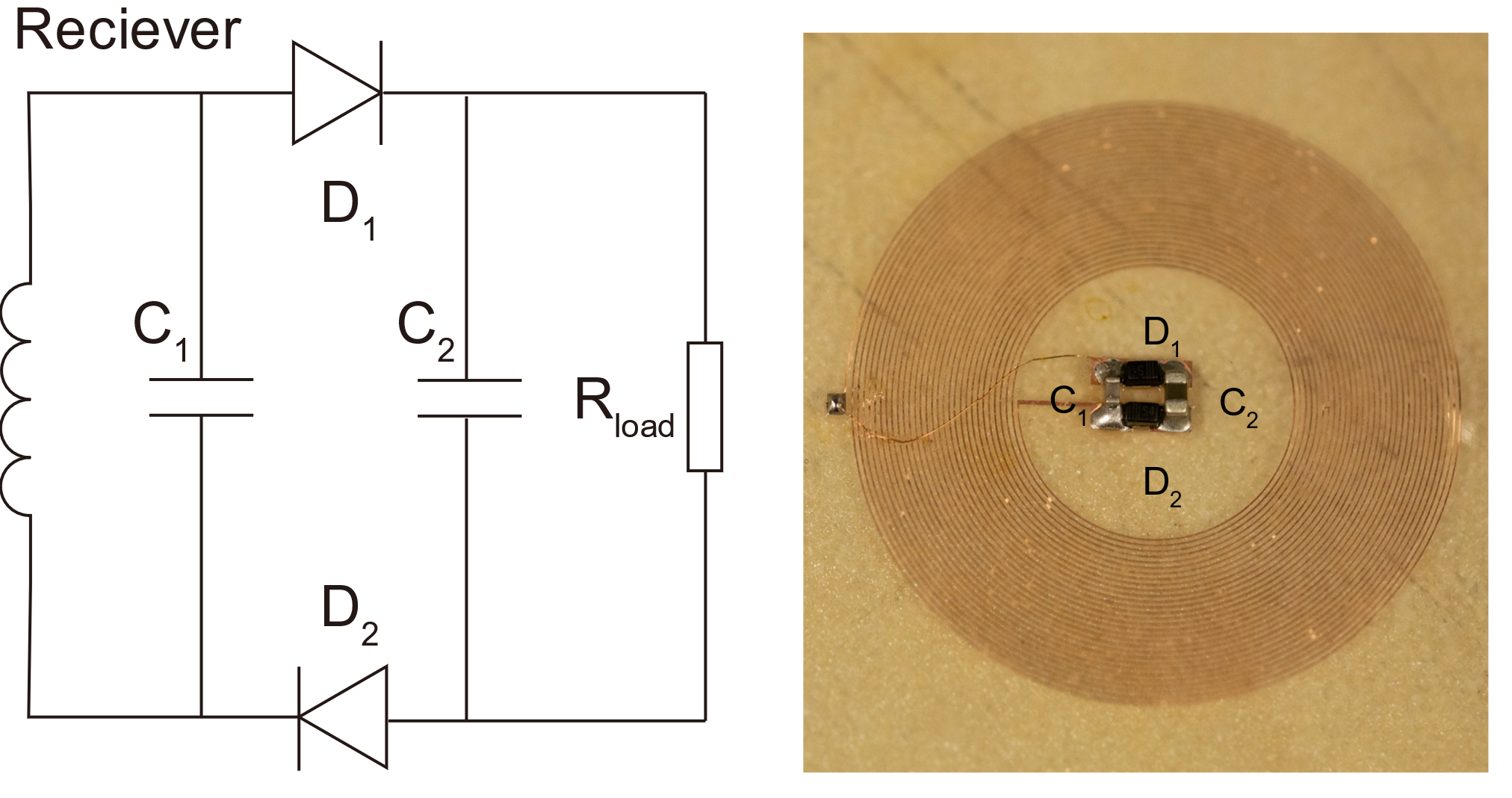

**Figure S8**. The circuit design (left), and the optical image (right) of the receiver coil.

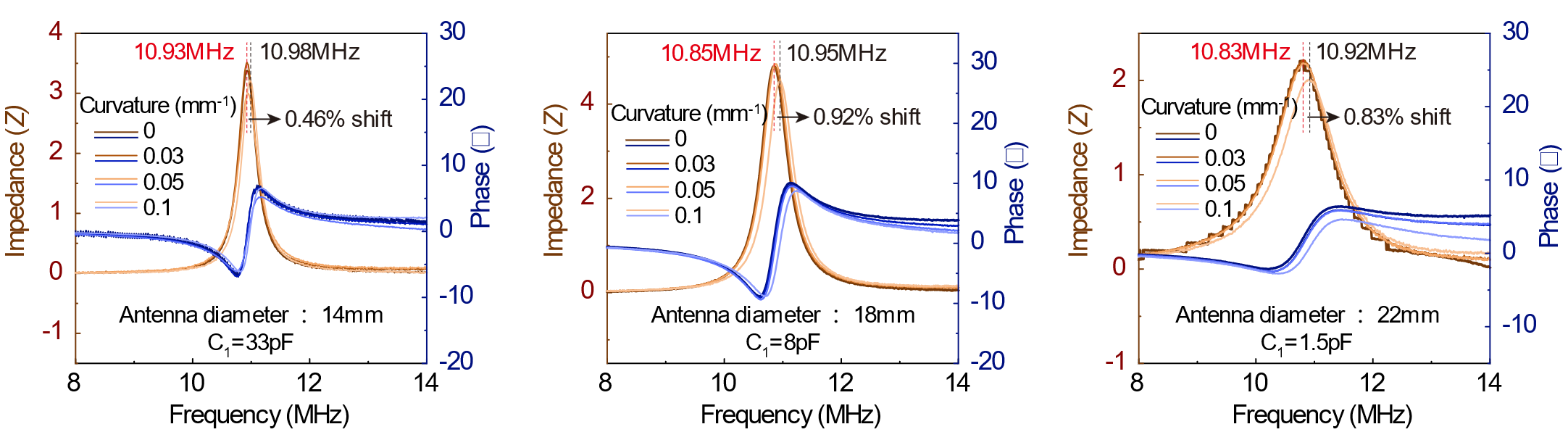

**Figure S9.** Resonant frequency characterizations for coils of different diameters (14, 18, and 22 mm) under different bending curvatures. The peak impedance indicates the resonant frequency for each condition.

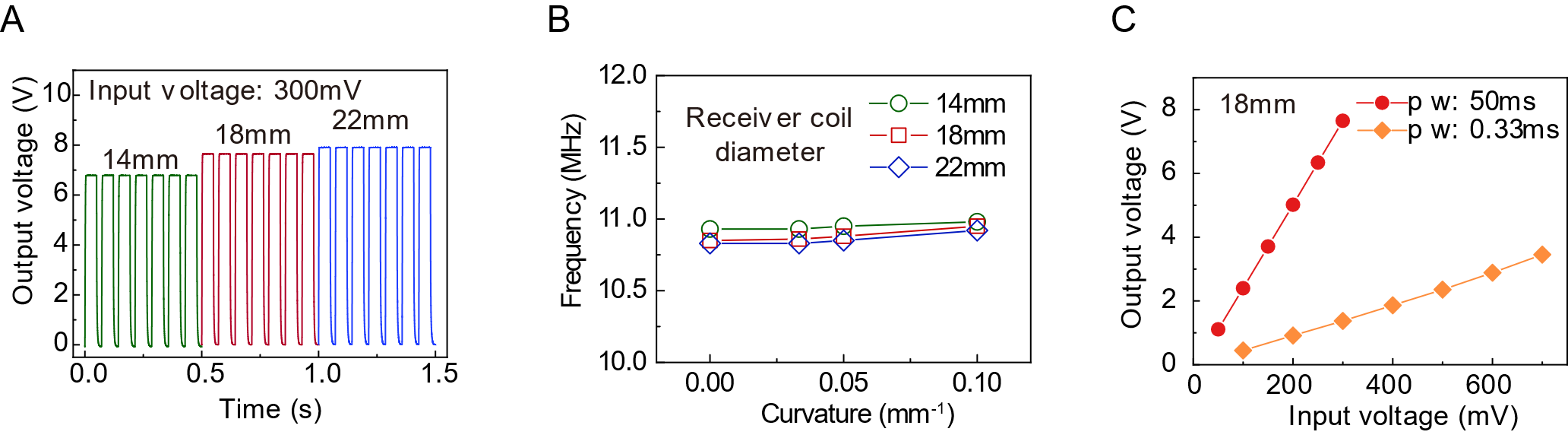

**Figure S10.** Characterizations of transmission properties. (A) Output voltage of the coil with different diameters (14, 18, and 22 mm) under the same pulse input. (B) The resonant frequency of receiver coils with different diameters (14, 18, and 22 mm) at different bending curvatures. (C) Output voltage as a function of input voltage with different pulse width (0.33 and 50 ms) as input pulses.

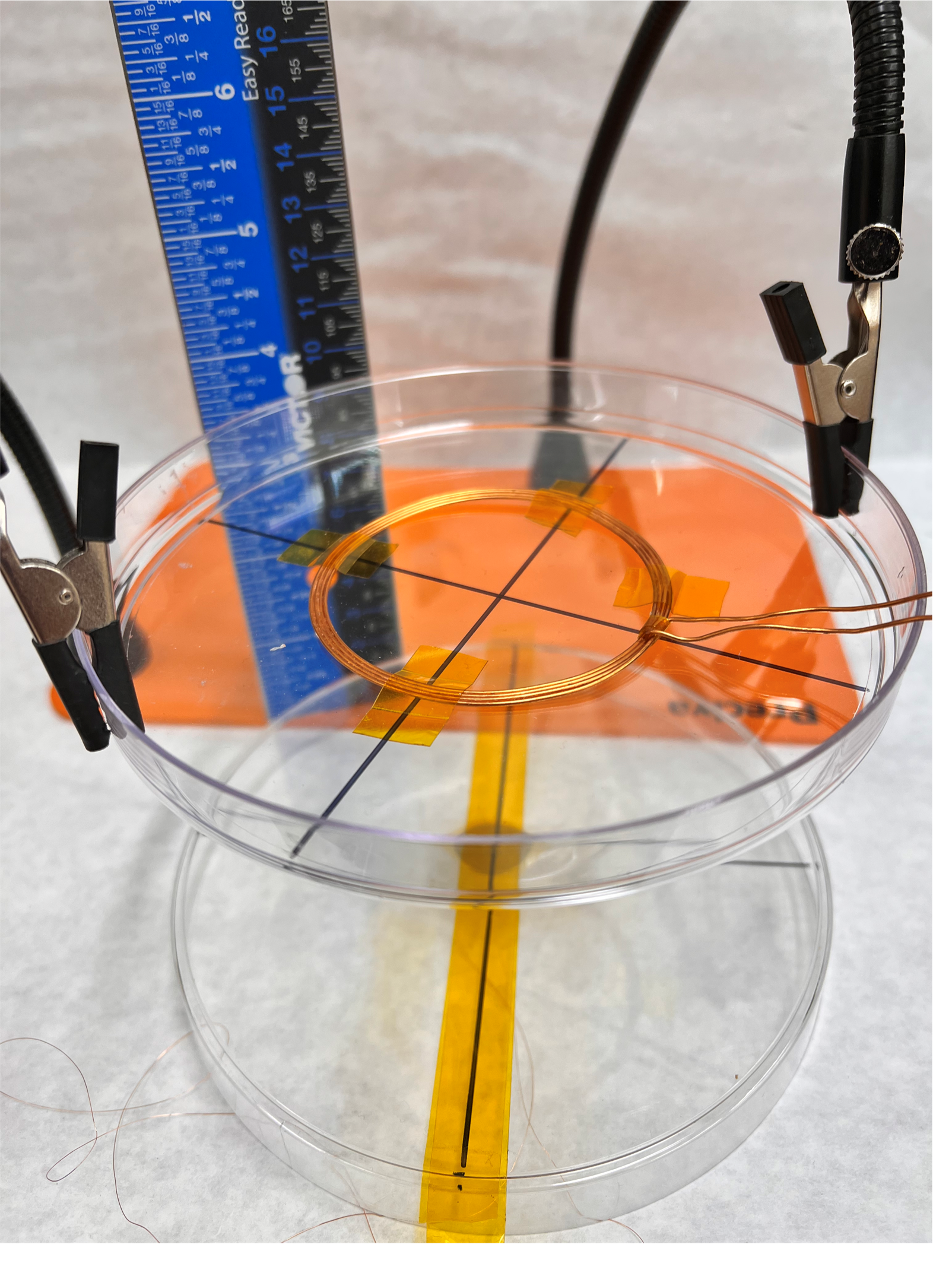

**Figure S11.** Optical image of in vitro coil characterization setup. Distance between the transmitter and receiver coil is measured using a ruler placed perpendicular to the two coils.

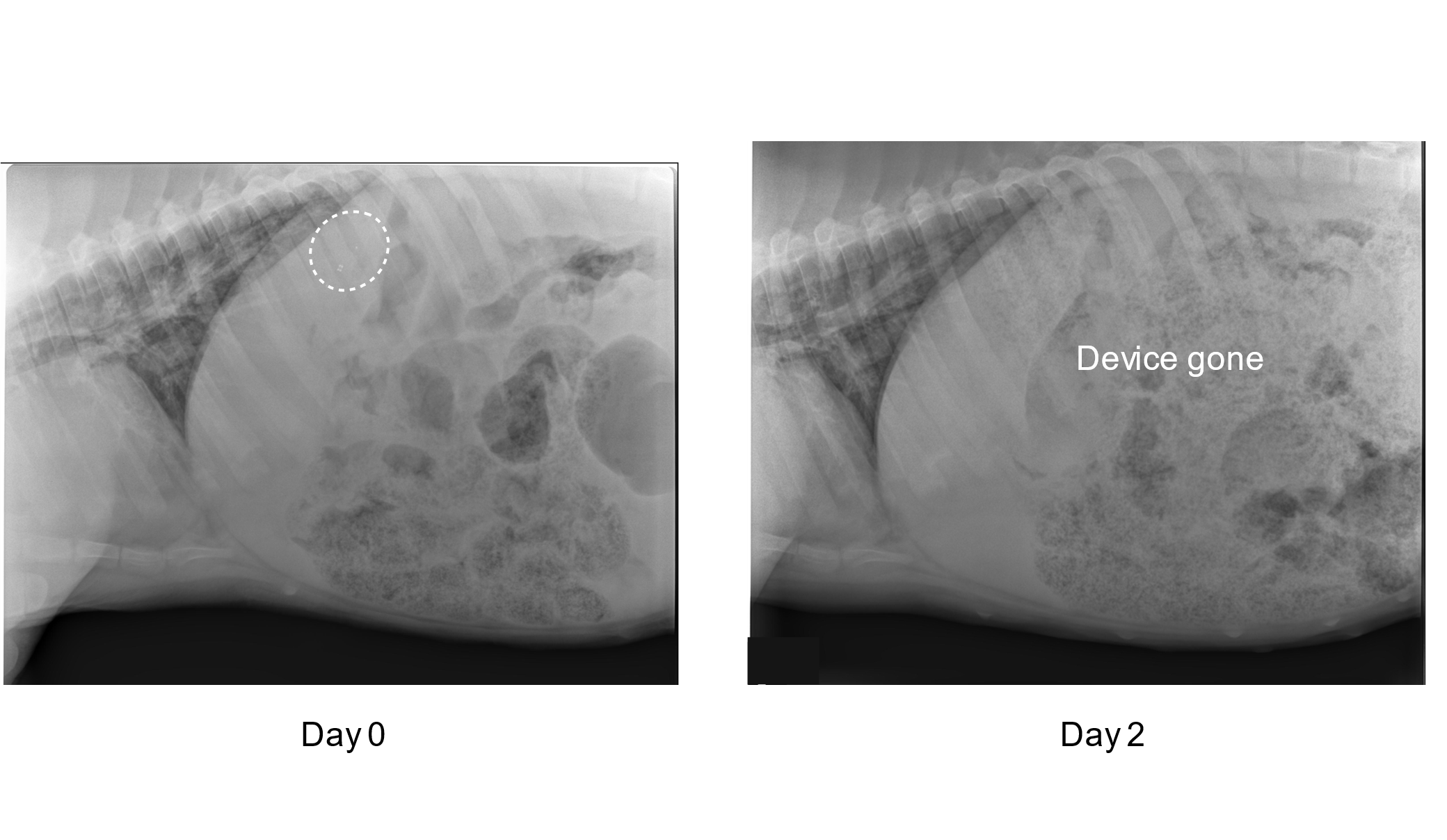

**Figure S12.** Radiographs of the ingested IngRI in the porcine stomach without the adhesive interface. On day 2, all devices safely exit the GI tract.
